# *Vibrio cholerae* phage ICP3 requires O1 antigen for infection

**DOI:** 10.1101/2023.01.31.526503

**Authors:** Drew A. Beckman, Christopher M. Waters

## Abstract

In its natural aquatic environment, the bacterial pathogen *Vibrio cholerae*, the causative agent of the enteric disease cholera, is in constant competition with bacterial viruses known as phages. Following ICP3 infection, *V. cholerae* cultures that exhibited phage killing always recovered overnight, and clones isolated from these regrowth populations exhibited complete resistance to subsequent infections. Whole genome sequencing of these resistant mutants revealed seven distinct mutations in genes encoding for enzymes involved in O1 antigen biosynthesis, demonstrating that the O1 antigen is a previously uncharacterized putative receptor of ICP3. To further elucidate the specificity of the resistance conferred by these mutations, they were challenged with the *V. cholerae*-specific phages ICP1 and ICP2. All seven O1 antigen mutants demonstrated pan-resistance to ICP1 but not ICP2, which utilizes the OmpU outer membrane protein as a receptor. We show that resistant mutations to ICP1 and ICP3 evolve at a significantly higher frequency than ICP2, but these mutations have a significant fitness tradeoff to *V. cholerae* and are unable to evolve in the presence of an antimicrobial that mimics host cell defensins.

## MAIN

### Introduction

The Gram-negative aquatic pathogen *Vibrio cholerae*, the causative agent of the severe enteric diarrheal disease cholera, is primarily found in the saline conditions of estuaries and brackish water (1). Recent estimates from 2015 place the human burden of disease worldwide in the range of 1.3-4.0 million cases and approximately 95,000 deaths annually as a result of rapid dehydration (2). Countries in sub-Saharan Africa and southern Asia are most directly impacted by cholera, but natural and manmade humanitarian disasters have ignited outbreaks in Haiti and Iraq (3, 4). Cholera is undoubtedly worsened by poverty where water treatment systems are suboptimal or non-existent (5).

Within its environment, *V. cholerae* is preyed upon by bacterial viruses known as bacteriophages, or phages, and these parasites are significant drivers of *V. cholerae* evolution (6). Lytic phages commandeer the host cell’s machinery and replicate to high numbers, eventually lysing the host cell. This predation is a critical driver of co-evolution among *V. cholerae* bacteria and their respective phages, applying selective pressure for the evolution of phage defense mechanisms like abortive infection (7), mobile genetic elements (8), and toxin-antitoxin systems (9) in addition to counter-defense systems evolved by the phage (10). Despite this robust phage defense, there are three major *V. cholerae-*specific lytic phage families isolated from patient stool samples at the International Center for Diarrheal Disease Research, Bangladesh known as ICP1, ICP2, and ICP3 (11, 12). These three phages, while identified together, are quite diverse; ICP1 is a member of the *Myoviridae* family and has a genome 2-3 times larger than ICP2 and ICP3 which are members of the *Podoviridae* family (12). The ICP phages are the three largest of the nine major phage families that have been identified to infect *V. cholerae* (13).

Before phage can successfully infect host bacteria, they must first bind to a specific cell surface receptor that determines phage specificity and host range. The receptor targeted by ICP1 to mediate phage invasion and replication is the O1 antigen in the lipopolysaccharide (LPS) of the *V. cholerae* outer membrane (11, 12). Of the over 200 known serogroups, only O1 and O139 cause cholera with the former responsible for the greatest number of cases (11, 14, 15). The El Tor biotype of *V. cholerae,* which is responsible for the current seventh cholera pandemic, can be subdivided into serotypes based on methylation of the O1 antigen. The Ogawa serotype is categorized by an additional methyl group on the most distal perosamine monomer while this methyl is absent in the Inaba serotype (4, 14, 15). The principal receptor for ICP2 is the OmpU outer membrane porin (16). However, the receptor for ICP3 remains uncharacterized (3, 17). O1 antigen and OmpU are excellent targets for phage exploitation because of their key roles in human pathogenesis and antimicrobial resistance. For instance, OmpU is required for efficient colonization of hosts while O1 antigen deficient *V. cholerae* mutants are more sensitive to the antibiotic polymyxin B as well as the bactericidal/permeability increasing (BPI) antimicrobial peptide secreted by the human gut (18, 19).

Understanding the interaction between the ICP phages and *V. cholerae* is not only essential for characterizing the evolution and ecology of *V. cholerae* in its natural environments, but also for the effective development of new cholera treatment strategies. Phage therapy is emerging as an effective complement to antibiotic treatment, especially with the rise of antibiotic resistant strains (20–22). The phage tropism for its individual receptor limits the negative effects on the human host and their microbiome (23). Two phage therapy approaches are being developed to treat cholera including the Phi_1 phage (24) and a cocktail of ICP1, ICP2, and ICP3 (3). Therefore, identifying the receptors of these phages has important clinical ramifications.

Here, we used a forward genetics approach to identify a previously unknown putative ICP3 receptor by examining several *V. cholerae* escape mutants following ICP3 infection. Our results showed that O1 antigen biosynthesis enzymes were mutated in all ICP3 resistant strains, and these mutants exhibited pan-resistance to ICP1 and ICP3, but not ICP2. ICP1 and ICP3 resistant *V. cholerae* evolve more readily than ICP2 resistant cells in laboratory conditions, but resistance to ICP3 fails to evolve in conditions that mimic host infection.

## Results

### Isolation of ICP3 resistant *V. cholerae*

During experiments studying the interaction of ICP3 phage with two newly discovered *V. cholerae* phage defense systems (7, 9), we observed that after an initial sharp decrease in the culture optical density upon ICP3 addition to values nearly equivalent to blank media controls, the bacterial cultures always rebounded to densities resembling the uninfected cultures. To establish the growth kinetics of *V. cholerae* in liquid media infected with ICP3 phage, an overnight culture of WT cells was subcultured into ten replicates, five of which were subsequently infected with ICP3 during mid-exponential growth phase at a multiplicity of infection (MOI) of 1.0 and the other five at 0.1 (Fig. 1). The density of the cultures rapidly decreased to initial inoculum levels 0.75 hr or 1.0 hr after ICP3 addition with 1.0 MOI or 0.1 MOI, respectively (Fig. 1). However, the OD600 of these cultures began to rebound 2 to 3 hr following the maximal drop. 24 hours later, the infected and uninfected cultures exhibited nearly identical optical densities (Fig. 1).

**Figure 1:**
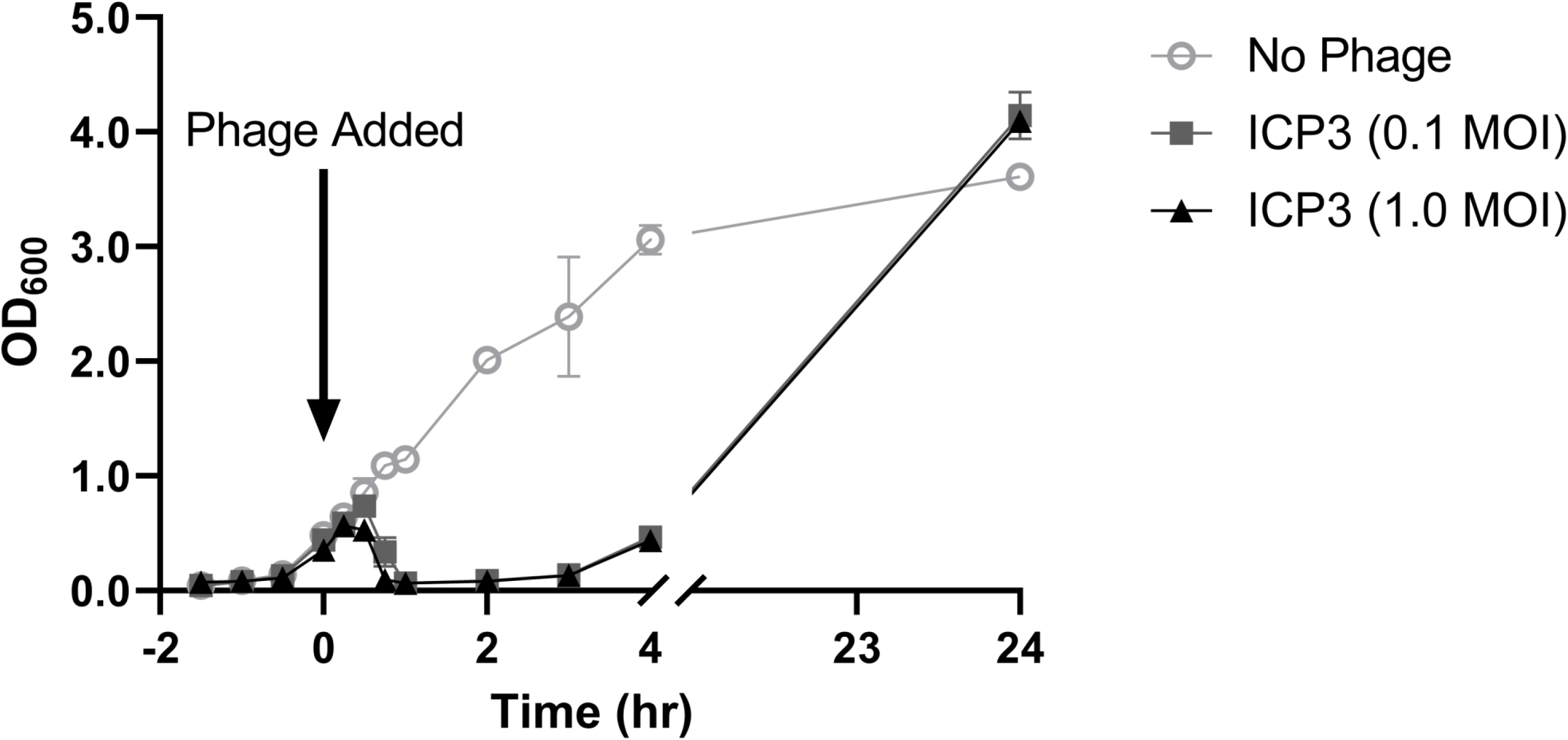
Infected *V. cholerae* cultures spontaneously recover over time. Growth curves of WT *V. cholerae* cultures challenged with ICP3 phage at varying multiplicities of infection (MOI). ICP3 phage, or SM buffer in the case of the no phage control, was added at 0 hr as indicated by the arrow. Data points are the mean of five biological replicates in the case of the phage groups (n = 5), and two in the case of the no phage control (n = 2). Error bars represent the standard deviation.

We hypothesized that the cells able to propagate in the phage-infected cultures were escape mutants resistant to ICP3. We isolated an individual clone from each of the ten independent overnight cultures that had rebounded from ICP3 infection. Each isolate was rechallenged with ICP3 at the original MOI of 1.0 for A-E (Fig. 2A) and 0.1 for F-J (Fig. 2B). Four of the five isolates infected with an MOI of 1.0 were resistant to ICP3 phage predation (Fig. 2A) while all five isolates challenged at an MOI of 0.1 were able to withstand ICP3 reinfection (Fig. 2B).

**Figure 2:**
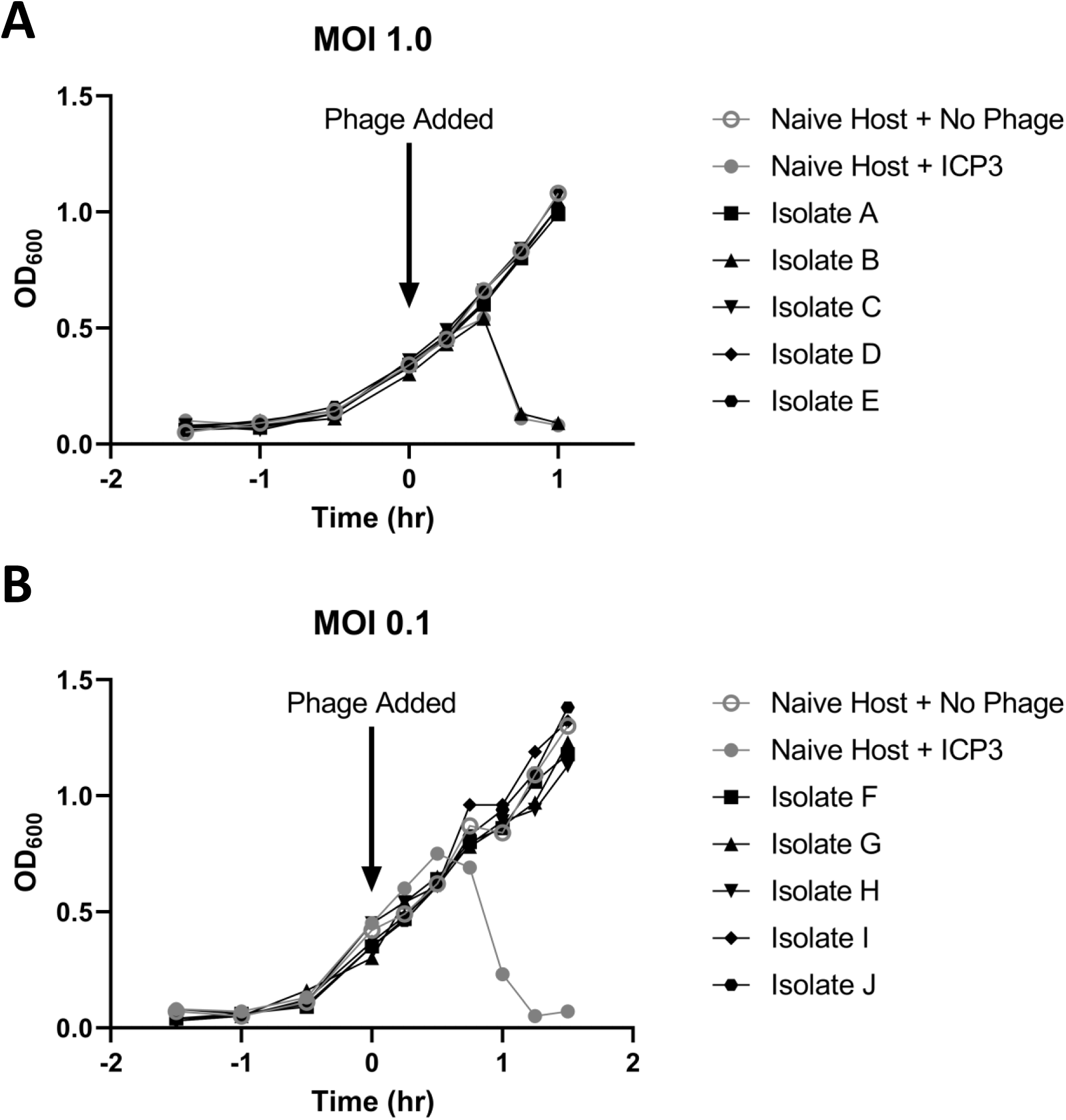
Recovered isolates demonstrate resistance to ICP3 phage in liquid culture. From the recovery pools of cultures that rebounded from ICP3 phage infection, individual isolates were obtained and rechallenged in a similar manner. They were either infected at an MOI of 1.0 (A) or 0.1 (B) based on their original recovery pool. ICP3 phage, or SM buffer in the case of the no phage control, was added at 0 hr as indicated by the arrow. Each line represents an individual isolate from an individual recovery pool.

### Resistant mutations map to the antigen biosynthesis gene cluster

Whole genome sequencing of all nine resistant isolates identified that isolates A-I (excluding the non-resistant isolate B) shared the same 11 bp deletion in the *manB* gene while isolate J had an insertion in the *wbeU* gene (Table 1, Fig. 3). Both genes are part of the O1 antigen biosynthesis operons of *V. cholerae* (14, 25). Given that the identical mutation in *manB* was isolated from 8 independent cultures, we hypothesized this mutation was likely present in the original common starting overnight culture. Therefore, to isolate additional resistance mutations, five independent overnight cultures of WT *V. cholerae* were inoculated using individual colonies and subjected to ICP3 infection at an MOI of 1.0. All five cultures exhibited regrowth after the initial population decline. Individual clones from these five populations that were resistant to ICP3 rechallenge were isolated and their genomes were sequenced (isolates K-O). In each isolate, there was only one unique mutation, all of which mapped to the O1 antigen biosynthesis pathway (Table 1, Fig. 3). Importantly, no other resistant mutations outside of the O1 biosynthesis genes were isolated. These seven unique ICP3 resistant mutants in the O1 antigen biosynthesis pathway were present in six different genes. Six of these mutations are likely null either by causing frameshift mutations (isolates A, J, L, N, and O) or a significant deletion (isolate K) while isolate M has a missense mutation in *gmd* (Table 1). Given the localization of these resistance-conferring mutations, we conclude that ICP3 requires the O1 antigen for infection of *V. cholerae*.

**Figure 3:**
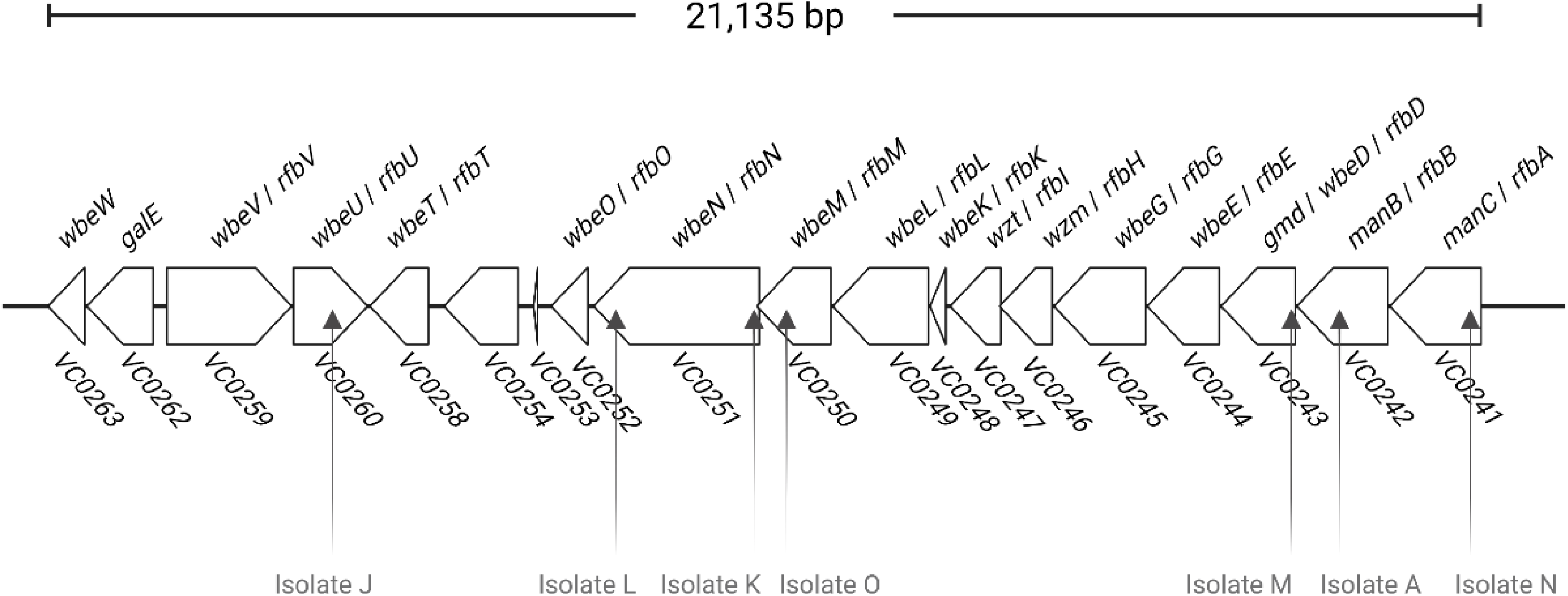
O1 antigen biosynthesis pathway genes indicating sites of mutations in resistant isolates. Diagram depicting the genes of the *wbe* locus (*rfb* locus) in O1 serotypes of *V. cholerae* strains. The block arrows of each gene point in the direction of transcription with gene sizes drawn to scale. Multiple names of each gene are given where applicable. The total length from the start codon of *VC0241* to the stop codon of *VC0263* is provided. Vertical arrows indicate the location of mutations in the individual isolates identified by whole genome sequencing.

**Table 1:** Whole Genome Sequencing of Pan-Resistant Isolates Reveals Mutations in the O-Antigen Biosynthesis Pathway.

| Isolate | Gene<sup>a</sup> | Position<sup>b</sup> | Location In Gene | Mutation | Protein Product<sup>c</sup> |
| --- | --- | --- | --- | --- | --- |
| A, C-I | *manB*, VC0242 | Chr I. 2,696,238 | 920-930/1392 nt | Δ11 bp | Phosphomannomutase |
| J | *wbeU*, *rfbU*, VC0259 | Chr I. 2,681,562 | 571/1113 nt | +G | Glycosyltransferase / LPS Biosynthesis Protein |
| K | *wbeN*, *rfbN*, VC0251 | Chr I. 2,687,710 | 78-95/2478 nt | Δ18 bp | Acyl-CoA Reductase / Acyl Protein Synthase |
| L | *wbeN*, *rfbN*, VC0251 | Chr I. 2,685,438 | 2367/2478 nt | +5 bp | Acyl-CoA Reductase / Acyl Protein Synthase |
| M* | *gmd*, VC0243 | Chr I. 2,695,683 | 91/1122 nt | G→A (H31Y) | GDP-Mannose 4,6-Dehydratase / Oxidoreductase |
| N | ***manC***, VC0241 | Chr I. 2,698,253 | 315/1398 nt | ΔC | Mannose-1-Phosphate<br>Guanylyltransferase / Mannose-6-Phosphate Isomerase |
| O | ***wbeM***, VC0250 | Chr I. 2,688,092 | 828/1125 nt | +A | Iron-containing Alcohol<br>Dehydrogenase |

<sup>a</sup>Gene names were determined as part of the *wbe* cluster (14, 15, 25) and the *rfb* cluster (28, 36).
<sup>b</sup>NZ\_CP046844 (Chr I) and NZ\_CP046845 (Chr II) used as reference genomes.
<sup>c</sup>Putative enzyme functions based on these reference genomes as well as Pombo *et al*, 2022 where applicable (37).
\*Demonstrated partial resistance in solid culture.

### ICP3 resistant mutants are also resistant to ICP1 but not ICP2 infection

To investigate if these seven ICP3 resistant mutants demonstrated cross-resistance to other ICP phages, the seven resistant mutants were tested in an efficiency of plaquing (EOP) assay for the ability to form plaques on a solid medium. An example of this plaquing assay for both the sensitive WT control and a representative resistant Isolate N is shown (Fig. 4A, B). Of the seven previously resistant *V. cholerae* mutants, six were completely resistant to ICP1 and ICP3 using EOP analysis showing no plaques at any dilution of phage tested (Figs. 4D, F). The WT and Isolate M exhibited identical numbers of plaques, suggesting this isolate does not exhibit equivalent resistance in plaque-based assays to that seen during liquid culture regrowth. However, the plaques of ICP1 or ICP3 that formed on Isolate M were significantly cloudier than the WT strain, suggesting this isolate has partial resistance to phage infection in these conditions (Fig. 4C). ICP2 efficiently infected all the ICP3 resistant mutants, supporting previous conclusions that ICP2 uses a different receptor, *ompU,* and not the O1 antigen (16).

**Figure 4:**
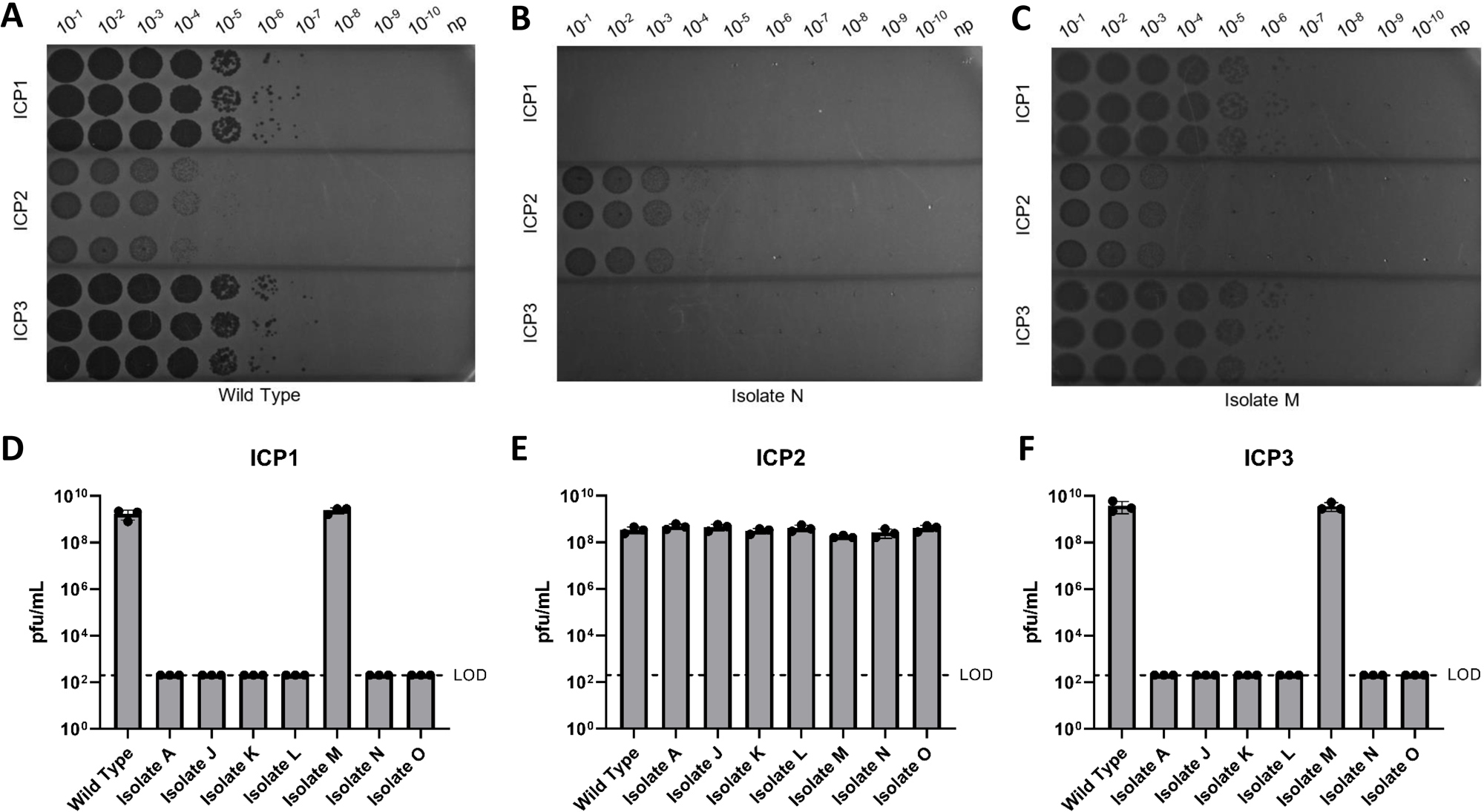
Efficiency of plaquing assay showing six isolates are pan-resistant to ICP1 and ICP3 phages. Seven isolates that were resistant to ICP3 in liquid culture were challenged with the three ICP phages on solid media. (A) A WT host is shown as a control. (B) Isolate N, a representative recovery isolate, is shown and is similar to Isolates A, J, K, L, and O. (C) Isolate M is shown depicting similar plaquing to WT but with a hazier phenotype. Dilutions used are shown across the top. np = no phage. (D-F) All three replicates for each phage are shown as dots and the bar indicates the mean. Error bars represent the standard deviation. The limit of detection (LOD) of this assay was 200 pfu/mL as indicated by the dashed line.

Given that all the ICP3 resistant mutants were able to regrow in liquid cultures when challenged with ICP3 yet Isolate M did not exhibit a decreased EOP compared with the WT, we performed quantitative liquid growth assays of the isolated resistance mutants challenged with ICP1 or ICP3 (Fig. 5). In these conditions, all the resistant mutants grew similar to the WT strain in the absence of phage (Fig. 5A) and were resistant to both ICP1 and ICP3 infection (Fig. 5B, C). Therefore, six of the seven mutations in the O1 antigen biosynthetic gene cluster exhibited resistance to both ICP1 and ICP3 in both EOP and liquid growth assays, while isolate M, with a missense mutation in *gmd,* was only resistant in liquid growth assays.

**Figure 5:**
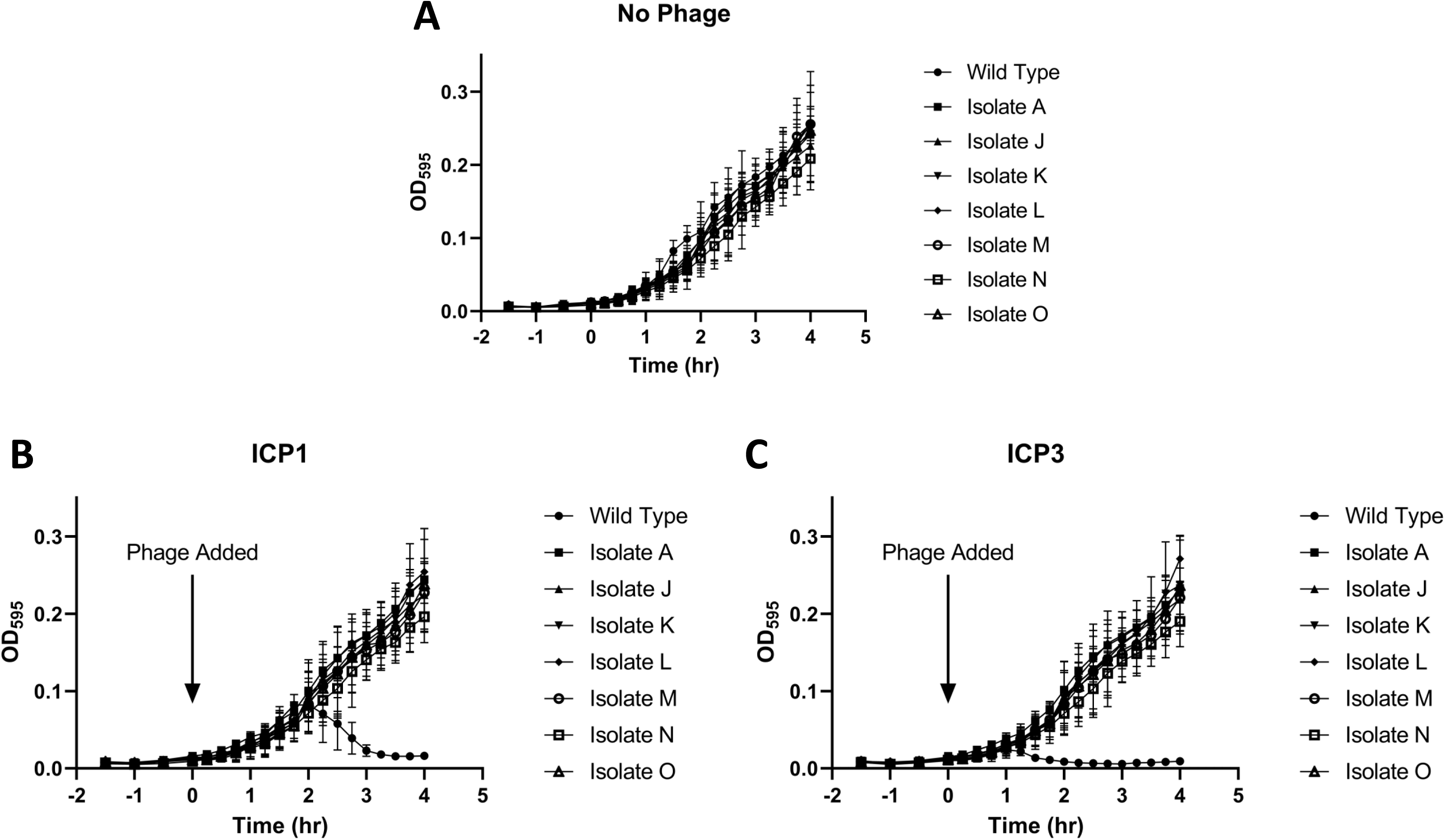
All ICP3 resistant mutants were pan-resistant to ICP1 in liquid culture. Growth curves for each ICP3 resistant mutant in the presence of no phage (A), ICP1 (B), and ICP3 (C) in liquid culture. Phage was added at 0 hr as indicated by the arrows. The average of three biological replicates is shown for each strain (n = 3). Error bars represent the standard deviation.

### Restoration of O1 antigen biosynthesis restores phage susceptibility

To ensure that phage resistance was due to the sequenced mutations, two representative isolates, A and J, were chosen and complemented by constructing inducible plasmids encoding *manB* and *wbeU*, respectively (Table I). Liquid infection assays compared the complemented strains and a deficient strain containing an empty vector. ICP1 or ICP3 phage were added at time 0 at a MOI of 1.0 (Fig. 6). In each case, the resistant isolates containing the complemented plasmid exhibited significant phage killing after 1 hr whereas the corresponding empty vector control strains were unimpacted by phage addition.

**Figure 6:**
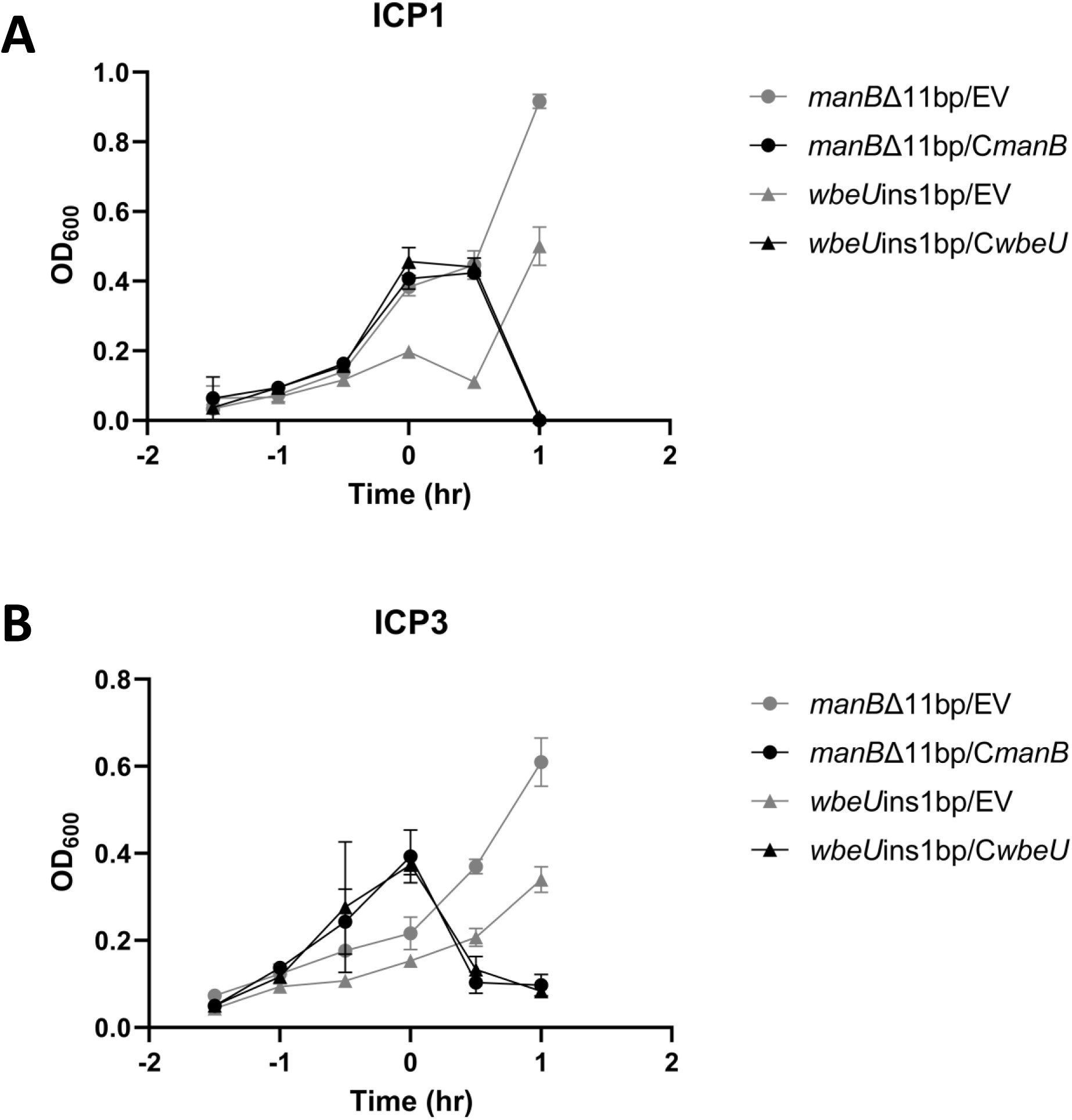
Complementation of mutated genes restores susceptibility phenotype. Mutant strains with inducible plasmids containing the intact *manB* gene (C*manB*), the intact *wbeU* gene (C*wbeU*), or anempty vector (EV) control were challenged with ICP1 (A) and ICP3 (B) at 0 hours. Growth curves were generated with time points every 0.5 hr. Results are shown as the mean of three replicates with error bars as the standard deviation.

### Resistance to ICP1 and ICP3 evolves more frequently than resistance to ICP2

These results and previous publications demonstrate that the target region for the evolution of mutants resistant to ICP1 and ICP3 is the O1 antigen biosynthetic cluster which spans a rather large 21,135 bp (12, 14). However, ICP2 is only known to utilize OmpU as a surface receptor, which is encoded by 1,026 bp, a much smaller target (16). We therefore postulated that the emergence of resistance to ICP1 and ICP3 in laboratory conditions would be significantly greater than to ICP2. To test this hypothesis, we performed a Luria-Delbrück fluctuation experiment with ICP1, ICP2, and ICP3 (26). For these experiments, WT *V. cholerae* was mixed with a large excess of phage on the surface of an agar plate, and the resulting phage resistant colonies were counted. Indeed, significantly higher numbers of resistant colonies were observed for ICP1 and ICP3 compared with ICP2 (Fig. 7). Given that *V. cholerae* can more easily evolve resistance to phages that target the O1 antigen in laboratory conditions, we next considered how such an infection strategy is evolutionary stable over time in the natural environments that *V. cholerae* occupies.

**Figure 7:**
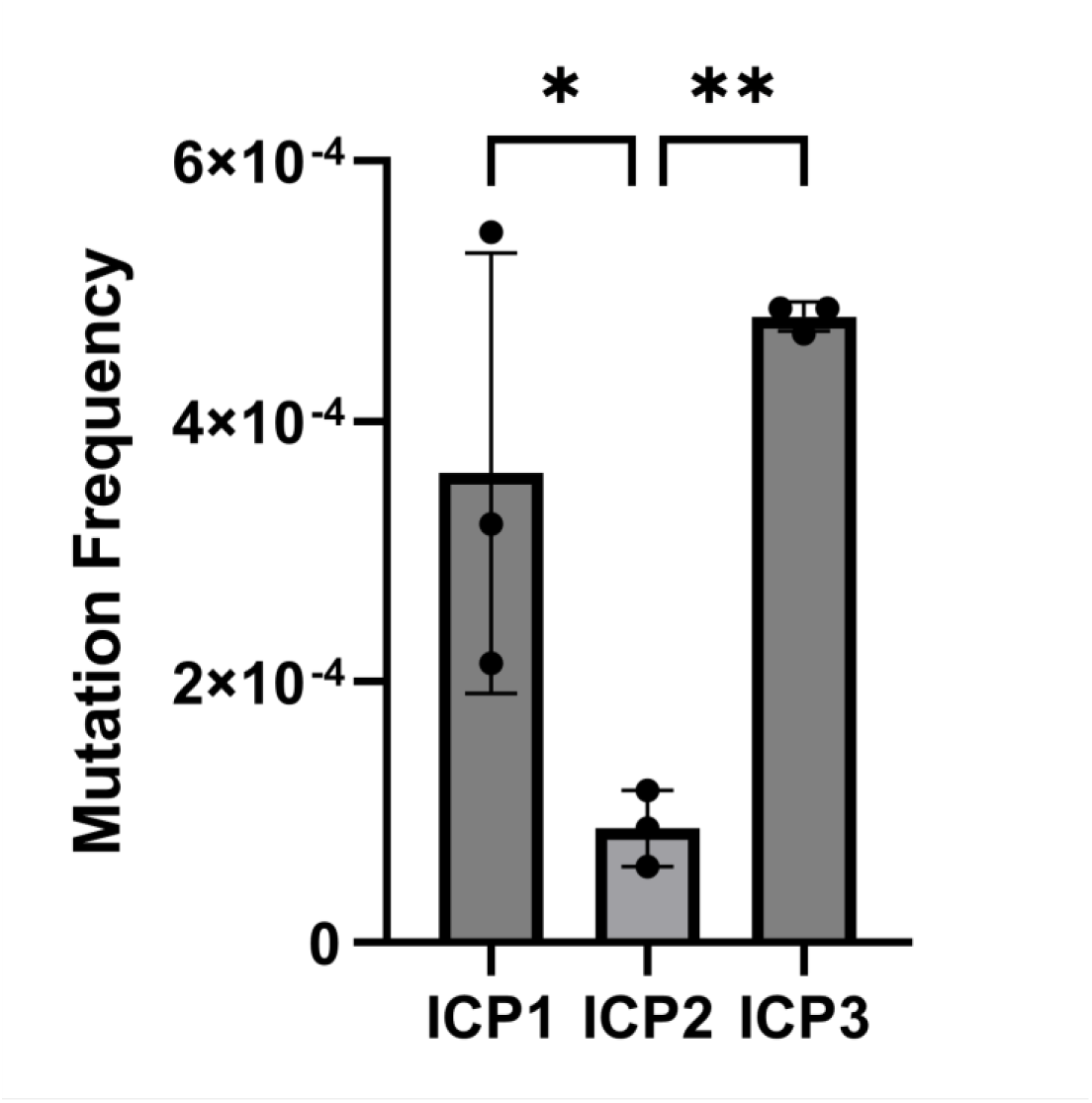
*V. cholerae* more readily evolves resistance to ICP1 and ICP3. A Luria-Delbrück fluctuation test was performed (n=3) using WT *V. cholerae* and ICP1, ICP2, or ICP3 as described in the materials and methods. The bars indicate the average mutational frequency, the error bars depict the standard deviation, and statistical significance was determined using a Two-Way ANOVA with Tukey’s post hoc multiple comparisons tests between each phage group, (*)p < 0.05, (**)p < 0.01.

### Enhanced sensitivity to polymyxin B decreases the evolution of ICP3 resistance

In Gram-negative bacteria, the O antigen provides an outermost protective layer to various stresses and antimicrobials (18). Previous O1 biosynthesis mutants that were resistant to ICP1 infection exhibited increased sensitivity to the antimicrobial polymyxin B (14). We therefore performed a minimum inhibitory concentration (MIC) assay to test if the seven O1 antigen ICP3 resistant mutants were likewise sensitive to polymyxin B (Fig. 8A, Table 2). Indeed, all seven mutants exhibited increased sensitivity to polymyxin B relative to the WT strain to varying degrees with some isolates (Isolate A and J) being approximately 30X more sensitive (Table 2). Isolate M, which was less resistant to ICP1 and ICP3 infections in EOP assays, most resembled the WT strain and was the only resistant mutant that was not significantly different than the WT. These results provide further evidence that these mutations disrupt O1 antigen biosynthesis although the extent of the results are variable depending on the specific mutation.

**Figure 8:**
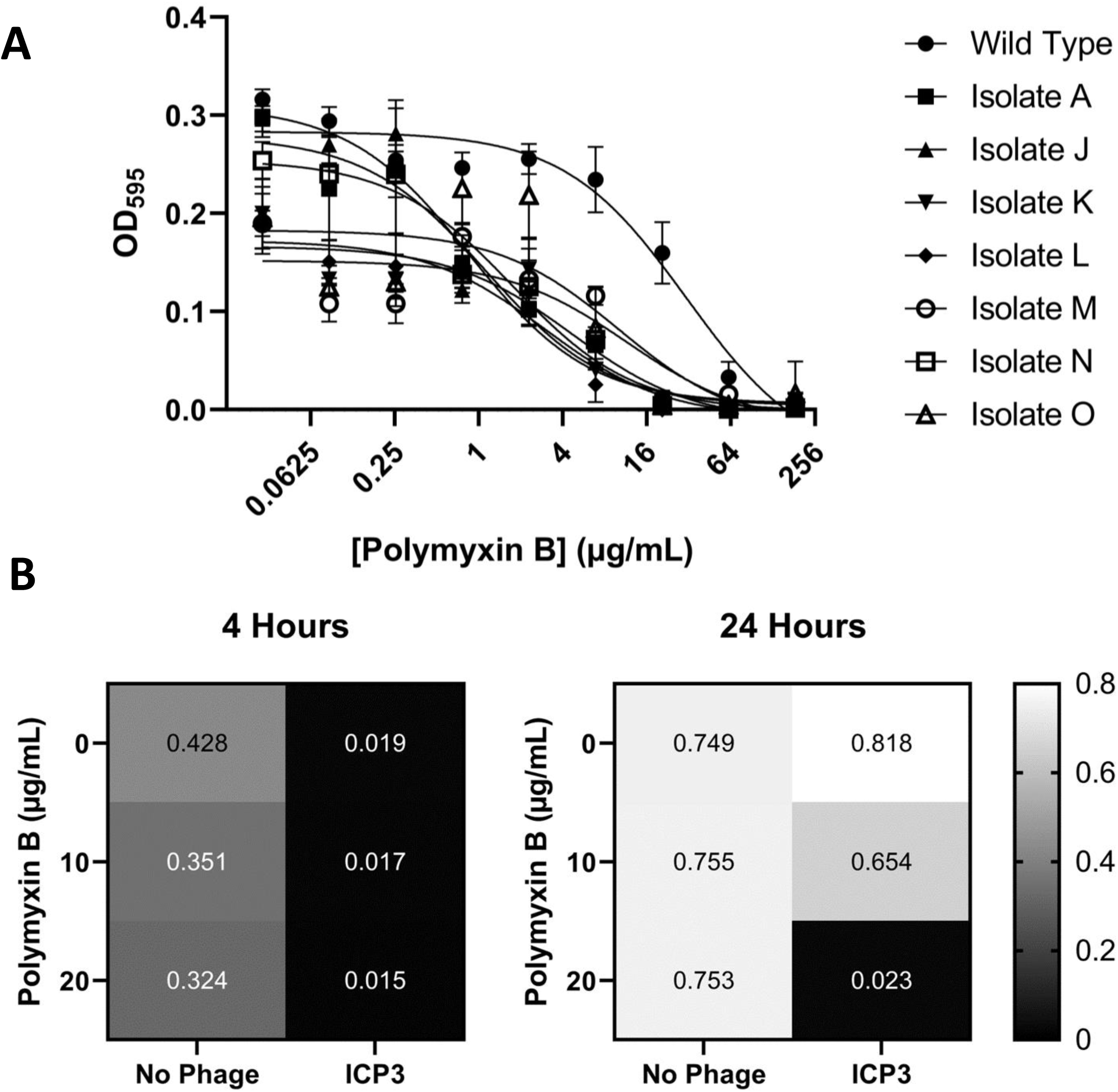
ICP3 resistance enhances sensitivity to polymyxin B. (A) A minimum inhibitory concentration curve was generated against the antibiotic Polymyxin B for all seven resistant isolates and WT. Lines of best fit were calculated using non-linear regression models to determine IC50. Each condition includes six biological replicates, and error bars depict the standard deviation. (B) The average OD595 of twenty independent wells for each condition at 4 and 24 hours is shown. ICP3 phage were added at 1-hour post-inoculation.

**Table 2:**
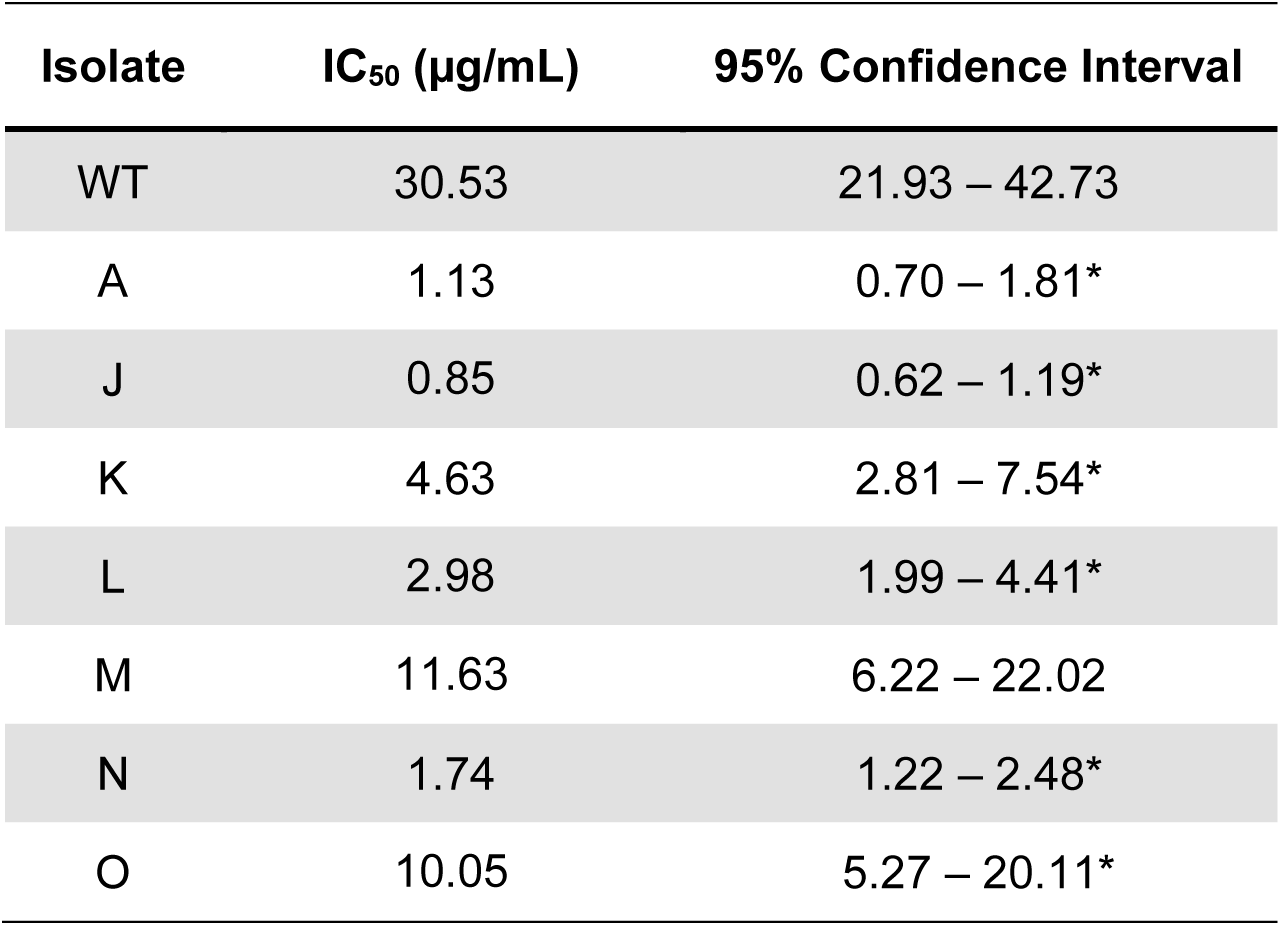
Half-Maximal Inhibitory Concentrations of Phage Resistant Mutants to Polymyxin B.

| Isolate | IC<sub>50</sub> (μg/mL) | 95% Confidence Interval |
| --- | --- | --- |
| WT | 30.53 | 21.93 – 42.73 |
| A | 1.13 | 0.70 – 1.81* |
| J | 0.85 | 0.62 – 1.19* |
| K | 4.63 | 2.81 – 7.54* |
| L | 2.98 | 1.99 – 4.41* |
| M | 11.63 | 6.22 – 22.02 |
| N | 1.74 | 1.22 – 2.48* |
| O | 10.05 | 5.27 – 20.11* |

Polymyxin B is a peptide antibiotic that mimics host cell cationic peptide defensins that target *V. cholerae* (19). Therefore, we hypothesized that selecting for ICP3 resistance in the presence of polymyxin B mimics an important selective condition of the host environment, and ICP3 resistance would be less likely to evolve in this condition. To test this, we quantified the culture rebound in 20 independent cultures of WT *V. cholerae* treated with no polymyxin B or the antibiotic at 10 and 20 µg/mL. Importantly, 20 µμg/mL is a concentration of polymyxin B that is below the MIC of the WT but above the MIC of all seven ICP3 resistant mutants we had isolated (Table 2). At four hours after ICP3 addition, all cultures treated with polymyxin B had low OD595 readings, indicating the phage had nearly eradicated the cultures. Uninfected cultures all robustly grew although there was a clear decrease in the OD595 with the addition of 10 and 20 µg/mL polymyxin B (Fig. 8B). By 24 hours, all 20 ICP3 infected cultures in the absence of polymyxin B or at a concentration of 10 µg/mL had rebounded, showing significant growth. However, none of the 20 cultures treated with ICP3 in the presence of 20 µg/mL polymyxin B rebounded and they all exhibited low OD595 values (Fig. 8B). Importantly, all cultures not exposed to phage grew to similar levels, indicating that the WT strain can grow in the concentrations of polymyxin B used. Thus, the evolution of resistance to ICP3 is inhibited by cationic antimicrobial peptides.

### Discussion

Our finding that *V. cholerae* cultures rebound after ICP3 infection through enrichment of escape mutants is consistent with the results of previous studies (3, 16, 27). Interestingly, all of these resistant mutants had defects in genes coding for enzymes that synthesize the O1 antigen. A putative pathway for the biosynthesis of the O1 antigen has been proposed by Stroeher *et al*, 1995 (28) with modifications by Chatterjee & Chaudhuri, 2004 (25) and Seed *et al*, 2012 (14). Three of the mutant strains in Table 1 had alterations to genes encoding perosamine synthesis enzymes. These include a frameshift deletion in the phosphomannomutase *manB* (Isolate A), a missense mutation in the GDP-mannose dehydratase *gmd* (Isolate M), and a frameshift deletion in the mannose-1 phosphate guanylyltransferase *manC* (Isolate N) (Table 1). Mutations that alter the function of these proteins prevent synthesis of the perosamine monomer in O1 antigen from being produced. *manB* mutations have been linked with *V. cholerae* ICP1 resistance, but not with resistance to ICP3 (27). Likewise, two mutations, one in-frame deletion (Isolate K) and one frameshift deletion (Isolate L), in the *wbeN* acyl-CoA reductase gene conferred pan-resistance to ICP1 and ICP3 phages (Table 1). The final isolates had a mutation in *wbeU* that is associated with LPS biosynthesis and *wbeM*, whose function in O1 antigen biosynthesis is unclear.

Given that the receptor for ICP1 has been previously identified as the O1 antigen, it is not surprising to see resistance among the isolates generated in this study. Therefore, both ICP1 and ICP3 require the O1 antigen for infection. However, ICP2 utilizes a different infection strategy. OmpU was previously identified as a receptor for ICP2 (16), and it was recently determined that the ICP2 tail protein Gp23 interacts with OmpU (29). OmpU was identified as the receptor of ICP2 from clinical stool samples by analyzing *V. cholerae* resistant to ICP2. These resistant cells had mutations in *ompU* or *toxR,* a transcriptional activator of *ompU* (16). The mutational target size for resistance to ICP1 and ICP3 is potentially 20 times larger than the target for ICP2, and we experimentally demonstrate that such a difference leads to more frequent evolution of *V. cholerae* resistance to ICP1 and ICP3 versus ICP2 in laboratory conditions. It should be noted that mutations in *toxR,* which is required for *ompU* expression, may also provide resistance to ICP2 and thus the target size for resistance mutations might be larger (3).

This study is the first to conclusively report the O1 antigen as a putative receptor for ICP3. A study exploring VP4 phage which is genetically similar to ICP3 and grouped as part of the ICP3 cluster, described its receptor in this same locus, but VP4 was not identified as ICP3 (13, 15). This study found mutations in *manB*, *manC*, *wbeU*, *wbeN*, and *wbeG*, which reinforce our findings here. Likewise, whole genome sequencing analysis of ICP3 resistant mutants isolated from infected mice showed mutations in the O1 biosynthesis genes, although these mutations were not further examined (3). Interestingly, some of these mutations showed differential resistance or sensitivity to ICP1, unlike those we isolated here which were all resistant to both phages. Our isolation of seven independent mutations in O1 antigen biosynthesis and no other putative receptors may suggest that the O1 antigen may be the sole receptor for ICP3. However, given that the target region for disruption of O1 antigen biosynthesis is large (Fig. 3), further studies with more saturating mutagenesis are required to conclusively test this hypothesis. As selection for O1 antigen resistant mutations is greatly decreased in the presence of 20 µg/mL polymyxin B (Fig. 8), and such an experimental condition would be useful to search for other factors that may be necessary for ICP3 infection.

The results here raise an important question as to why these O1 antigen deficient *V. cholerae* lineages have not dominated environmentally since they provide pan-resistance to ICP1 and ICP3 infections. Since LPS operates as a virulence factor within the human host, it is important for survival and replication in that niche (12). LPS mutants exhibit decreased competitiveness and ability to colonize the human gut compared to their wild type counterparts (30). There is additional evidence that the LPS protects against acid exposure which may be typical during human colonization (30). These collective results suggest that emergence of O1 antigen null mutants that provide resistance to ICP3 is not evolutionary favored due to their significant fitness cost. This finding is consistent with previous studies that have similarly concluded that although isolation of phage receptor mutants is common during selection for phage resistance in laboratory conditions, such mutants are rarely seen in natural populations (31). In support of this, twenty independent cultures of *V. cholerae* infected with ICP3 failed to evolve resistance in the presence of 20 µg/mL polymyxin B, an experimental condition that more closely mimics the host environment.

The E7946 El Tor strain encodes homopolymer nucleotide tracts in the genes *wbeL* and *manA,* which are required for O1 antigen biosynthesis, that mediate phase variable expression of the O1 antigen leading to ICP1 phage resistance (14). It was postulated that such a system could lead to a strategy in which a segment of the population evolves resistance to ICP1 phage infection while maintaining the genetic potential to restore O1 for infection of the host. We did not isolate any mutations in *manA* or *wbeL*, and an analysis of the sequence of genes in the El Tor strain used here, C6706, showed that the polyA(A9) tracts were conserved in *manA* but not in *wbeL.* This suggests that the loss of *manA* is not favored upon ICP3 selection.

As phages use cell surface proteins and molecules as receptors, an emerging concept is that resistance to phage infection may have the evolutionary tradeoff of antibiotic sensitivity. Such a tradeoff has been demonstrated in the *E. coli* phage U136B which uses the antibiotic efflux protein TolC as a receptor, as mutations that confer resistant to phage infection lead to increased antibiotic sensitivity (32). Evolving to recognize the O1 antigen as a receptor by ICP1 and ICP3 is a natural example of such a concept as any *V. cholerae* that gain resistance through loss of O1 have the tradeoff of loss of virulence and more sensitivity to host antimicrobials that resemble polymyxin B. Therefore, evolution to use the O1 antigen for infection by ICP1 and ICP3 has contributed to the evolutionary success of these lytic phages.

## MATERIALS AND METHODS

### Bacterial culturing and molecular biology

The WT culture is El Tor *V. cholerae* C6706:str2 (33). All cultures were grown in glass test tubes (18 x 150 mm, Kimax) containing Luria-Bertani broth at 35°C with shaking, unless otherwise stated. Selection of plasmids utilized kanamycin at 100 µg/mL. The *manB* (pDAB1) and *wbeU* (pDAB2) complementation plasmids were generated using Gibson cloning. The gene inserts were generated from C6706:str2 with GoTaq 2x MasterMix using the primers (overhang, <u>RBS</u>, INSERT): 5’-acagcctcgacaggcctagg<u>aggagctaaggaagctaaa</u>GTGAAAGAGTTAACTTGTTTT-3’ and 5’-gcttgctcaatcaatcaccgTTAAATATCCAATTTCTTAATTAATTTAGTAAG-3’ (*manB*) 5’-acagcctcgacaggcctagg<u>aggagctaaggaagctaaa</u>ATGCCATGGAAGACCTAC-3’ and 5’-gcttgctcaatcaatcaccgTCAACAGACATTTCCGAAG3’ (*wbeU*). These inserts were Gibson (New England Biolabs) cloned into an EcoRI/BamHI digested pEVS143 plasmid (34) using standard methodologies followed by Sanger sequencing to confirm the proper clone was generated. The empty vector and plasmids were conjugated from the *Escherichia coli* diaminopimelic acid auxotrophic strain BW29427 (K. Datsenko and B.L. Wanner, unpublished) into *V. cholerae* strains by mixing 25 µL of each culture overnight on an agar plate followed by selection on LB plates with kanamycin.

### Liquid Infection Assays

30 µL of overnight cultures were back diluted into 3 mL of LB and allowed to grow until OD600 (0.20-0.50). Each culture was then infected with ICP1 (ICP1_2011_A - MH310933), ICP2 (ICP2_2013_A_HAITI - KM224879), ICP3 (ICP3_2007_A - HQ641344), or SM buffer (50 mM Tris-Cl + 100 mM NaCl + 8 mM MgSO4). Phage stocks were maintained in filtered LB medium with trace chloroform. The amount of phage to add was determined by converting the optical density (OD) of the culture into an estimated viable cell count in *cfu/mL* using a quadratic regression equation generated from a previous growth curve: *cfu/mL* = [1.4226 x 10^9^ x (*OD*)^2^] + [4.2981 x 10^8^ x *OD*] – 2.2521 x 10^6^. Following infection, the OD600 of cultures was remeasured every 0.25 hr. For the recovery experiments, cultures were allowed to recover for 24 hours. The recovery pools were collected and frozen in 20% glycerol at -80°C. The recovery pools were then struck onto LB agar plates and individual isolates were used for reinfection. The same infection procedure was followed for the reinfection assays. For the liquid phage infection assays in Fig. 4, three independent overnight cultures of the WT or seven phage resistant isolates grown in 2 mL of LB were back diluted 1/100 in fresh LB and 100 µL was added to four wells into a clear 96-well plate (Costar). The plate was incubated at 35°C for 1.5 hours and to one well 1 µL of the ICP1 or ICP3 phage stock was added. This led to MOIs of 0.5 and 0.36 for ICP1 and ICP3, respectively. The OD595 was monitored every 0.25 hours in an Envision plate reader. For the evolution of ICP3 resistance in polymyxin B in Fig. 8, the experiment was performed similarly with the exception that twenty independent overnight cultures of WT *V. cholerae* were inoculated in 150 µL of LB within a 96-well plate and grown overnight at 35°C in a humidity chamber without shaking. These cultures were then used to inoculate 40 wells of 150 µL of LB with either 0, 10, or 20 µg/mL polymyxin B with a pin transfer tool. After one hour of growth, 1 µL of ICP3 phage was added to half of the wells for each condition (n=20), which approximates to a MOI of 1.0. The plates were incubated at 35°C in a humidity chamber without shaking. The OD595 of all wells were measured at 4 and 24 hours using an Envision plate reader.

### Efficiency of Plaquing Assays

Cultures of WT and resistant strains were grown overnight in LB and 250 µL was inoculated into 14 mL warm MMB (LB + 0.5% agar + 5 mM MgCl2 + 5 mM CaCl2 + 0.1 mM MnCl2) at 60°C. The mixture was poured onto an empty plate (150 x 15 mm, Fisher Scientific) and allowed to solidify. Ten-fold serial dilutions of phage made in SM buffer were then spotted 5 µL in succession onto each strain along with one SM buffer spot as a negative control. This was performed in triplicate for each strain. Once plates were dried for about 20-30 minutes, they were allowed to incubate at 35°C overnight. Plaques were counted the next day to determine the efficiency of plaquing on each strain for each phage (ICP1, ICP2, and ICP3).

### Fluctuation assay

Three independent cultures of WT *V. cholerae* for each ICP phage were grown overnight in 3 mL of LB 35°C overnight with shaking. The following day, 1 mL of phage was evenly spread on an agar plate. All phages were diluted in LB to a titer of 6.0 x 10^7^ pfu/mL to ensure consistency between phage groups. Before the plate was dry, 1 µL of a 1/100 dilution of the overnight culture was also spread on the plate, resulting in a MOI of ∼100. Resistant colonies were quantified at 4 days excluding all colonies forming on the edges of the plates, as these were frequent and likely did not interact with the phage on the plate. The frequency of mutation was calculated by multiplying the number of mutant colonies by the dilution factor of 10^2^ and then dividing by the average cfu/mL (1.03 x 10^10^ cfu/mL) of all nine starting cultures to normalize dilution error. This was determined by spotting 5 µL serial dilutions of the overnight cultures on agar plates.

### Whole Genome Sequencing and Analysis

Genomic extraction for the WT C6706:str2 strain and each of the resistant mutants was performed using Wizard Genomic DNA Purification Kit (Promega). The concentration of DNA was quantified using Nanodrop spectrophotometer (ND-1000). Samples were then sequenced at SeqCenter (Pittsburgh, PA). Results were processed by aligning each sample to the reference chromosomes of *V. cholerae* (NZ_CP046844 and NZ_CP046845) using *breseq* (35). Any mutations found in the WT strain as well as our resistant mutants were removed from consideration.

### Polymyxin B minimum inhibitory concentration

Six independent overnight cultures of the WT and all seven resistant mutants were grown in 150 µL of LB in a 96-well plate overnight. Cultures were inoculated into 150 µL of LB with the indicated concentrations of antibiotics in a clear 96-well plate (Costar) using a pin transfer tool.These were then grown at 35°C without shaking in a humidity chamber for 6 hours (185-0.76, 0 µg/mL) or 4 hours (0.25-0.0282 µg/mL) at which time the untreated WT culture grew to an OD595 of 0.3. The OD595 was measured using an Envision plate reader.

## ACKNOWLEDGEMENTS

This research was supported and funded by NIH grants GM139537 and AI158433 to C.M.W. D.A.B. was supported by a Professorial Assistant fellowship from Michigan State University and the Norman E. & Hanna C. Kelker Research Scholarship and the Dietrich C. Bauer Scholarship from the Department of Microbiology and Molecular Genetics. We thank Andrew Camilli and Wai-Leung Ng at Tufts University for providing the ICP1 and ICP3 phage and Kim Seed for providing us with the ICP2 phage, additional *V. cholerae* strains, and critical input.

